# Comparative study of gamma radiation tolerance between desiccation-sensitive and desiccation-tolerant tardigrades

**DOI:** 10.1101/2024.06.26.600756

**Authors:** Tokiko Saigo, Katsuya Satoh, Takekazu Kunieda

## Abstract

Tardigrades are small metazoans renowned for their exceptional tolerance against various harsh environments in a dehydrated state. Some species exhibited an extraordinary tolerance against high-dose irradiation even in a hydrated state. Given that natural sources of high radiation are rare, the selective pressure to obtain such a high radiotolerance during evolution remains elusive. It has been postulated that high radiation tolerances could be derived from adaptation to dehydration, because both dehydration and radiation cause similar damage on biomolecules at least partly, e.g., DNA cleavage and oxidation of various biomolecules, and dehydration is a common environmental stress that terrestrial organisms should adapt to. Although tardigrades are known for high radiotolerance, the radiotolerance records have been reported only for desiccation-tolerant tardigrade species and nothing was known about the radio-tolerance in desiccation-sensitive tardigrade species. Hence, the relationship between desiccation-tolerance and radio-tolerance remained unexplored. To this end, we examined the radiotolerance of the desiccation-sensitive tardigrade, *Grevenius myrops* (formerly known as *Isohypsibius myrops*) in comparison to the well-characterized desiccation-tolerant tardigrade, *Ramazzottius varieornatus*. The median lethal dose (LD_50_) of *G. myrops* was approximately 2,240 Gy. This was much lower than those reported for desiccation tolerant eutardigrades. The effects of irradiation on the lifespan and the ovipositions were more severe in *G. myrops* compared to those in *R. varieornatus*. The present study provides the precise records on the radiotolerance of a desiccation-sensitive tardigrade and the current data supported the certain correlation between desiccation tolerance and radiotolerance at least in eutardigrades.

## INTRODUCTION

Tardigrades, also known as water bears, are aquatic micrometazoans which can be found in various habitats such as marine, freshwater, and limno-terrestrial ecosystems (Møbjerg *et al*., 2011). Some terrestrial tardigrade species can endure desiccation by reversibly entering a dehydrated state referred to as anhydrobiosis, and they can also tolerate high-dose irradiation, sometimes reaching more than 5,000 Gy which corresponds to approximately 1,000 times higher than the median lethal dose of humans (Jönsson *et al*., 2005, 2016; Horikawa *et al*., 2006, 2008; Beltrán-Pardo *et al*., 2015; see the review, Hashimoto and Kunieda, 2017; Jönsson, 2019). Given that natural sources of high radiation are rare, the evolutionary force to reinforce the radiotolerance of animals to so high-level observed in some tardigrades remains elusive. One possible explanation about this is that high radiotolerance is a derivative of high desiccation tolerance. Desiccation is apparently much more common environmental stress for terrestrial organisms to cope with, and both desiccation and radiation may cause similar damage to biomolecules and cells, partly through the common generation of reactive oxygen species. Hence, the idea is that the machinery evolved to cope with desiccation may help animals survive when exposed to high-dose irradiation. Indeed, there were several observations that the radiotolerant organisms tend to have high desiccation tolerance, e.g., in insects (Ryabova *et al*., 2017), bdelloid rotifers (Gladyshev *et al.,* 2008) and bacteria (Billi *et al*., 2000; Cox *et al*., 2005).

However, radiation tolerance in tardigrades has been examined with a focus on desiccation-tolerant species (Hashimoto and Kunieda, 2017; Jönsson, 2019). Almost no information is available on desiccation-sensitive tardigrade species. To examine the relationship between radiotolerance and desiccation tolerance in tardigrades, the radiotolerance data of desiccation-sensitive species is indispensable. To this end, we focused on a highly desiccation-sensitive eutardigrade, *Grevenius myrops* (formerly known as *Isohypsibius myrops*, updated by Gąsiorek *et al*., 2019; Isohypsibioidea, Parachela, Eutardigrada). The rearing system of this species was established and the strain termed Im1 derived from a single parthenogenetic individual has been maintained for more than 10 years in a laboratory. The strain was reported to be quite sensitive to desiccation as it cannot withstand even exposure to 98% relative humidity (%RH) for 2days or 94-95%RH for 1 day (Ito *et al*., 2016). In this study, we analyzed the radiation tolerance of desiccation-sensitive *G. myrops* by exposing them to various doses of gamma rays and reported the detailed data about the effects on animal activity, lifespan, ovipositions, and hatchability of laid eggs. Our data provide a valuable foundation to elucidate the relations and evolutionary history between radiation tolerance and desiccation tolerance.

## MATERIALS AND METHODS

### Animals

As a desiccation-sensitive tardigrade, we used the Im1 strain of a freshwater tardigrade, *G. myrops*, which was previously established from a single individual and exhibited no tolerance against desiccation (Ito *et al*., 2016). As a desiccation-tolerant tardigrade species, *R. varieornatus* (YOKOZUNA-1 strain) was used (Horikawa *et al*., 2008). Both tardigrades were reared in 1.2% agar plates (STAR Agar H-grade 01, Rikaken, Japan) overlaid with thin layer of water, essentially as described previously (Ito *et al*., 2016, Horikawa *et al*., 2008). For *G.* myrops, freshwater rotifers *Lecane inermis* and live chlorella suspension (Chlorella Industry) were added as a food once a week. For *R. varieornatus*, only live chlorella suspension was added as a food. Tardigrades were kept at 22–23 °C under ambient photoperiod and were transferred to new agar dishes approximately once a month.

### Gamma-irradiation

As a container, 0.2 mL PCR tubes were used (Fig. S1). In each tube, 30 µL of 1.5% agar was solidified at the bottom, and 19–20 adult tardigrades were transferred with 30 µL of sterilized Milli-Q water (Millipore) on the agar. Tardigrades were irradiated with ^60^Co gamma rays at room temperature for 60 minutes at the Food Irradiation Facility, Takasaki Institute for Advanced Quantum Science, QST (Gunma, Japan). The irradiation dose rate ranged from 8.3 to 116.7 Gy/h, which was regulated by adjusting the distance of the samples from the gamma ray source. Desiccation-sensitive *G. myrops* were irradiated with 6 doses: 0 Gy (as a non-irradiated group), 500 Gy, 1,000 Gy, 2,000 Gy, 3,000 Gy and 4,000 Gy. Desiccation-tolerant *R. varieornatus* was irradiated with 4 doses: 0 Gy, 3,000 Gy, 5,000 Gy and 7,000 Gy. All samples were irradiated simultaneously, and 4 biological replicates were examined for each condition.

### Observation of Activity and Oviposition

We used a deep glass blood reaction plate (Toshin Riko) to observe animals after irradiation. The bottom of each well (22 mm in diameter) was filled with 150 µL of 1.5% agar and overlaid with 150 µL of sterilized Milli-Q water. After irradiation, tardigrades were transferred to a glass dish without agar (Agar Scientific) and were washed well with sterilized Milli-Q water (Millipore). Then, they were transferred with 50 µL of sterilized Milli-Q to an agar-laid glass blood reaction plate and were reared with a food as described above. The plates were incubated in a tight box and sterilized Milli-Q water was added daily to each well to compensate for the evaporation. Tardigrades were transferred to a new agar well at 4 days, 12 days, and 20 days after irradiation. *G. myrops* was fed every 2–4 days, whereas *R. varieornatus* was fed when it was transferred to a new agar well. We daily examined the number of active animals, egg laying and hatchability of laid eggs till 6 days after irradiation, and thereafter, every 2 days till 28 days post-irradiation. Animals moving their legs or internal organs were regarded as “active” animals. The median lethal dose (LD_50_) was calculated by fitting to probit model using the statistical software R.

### Observation of eggs laid by irradiated adults

The oviposition of irradiated adults was examined till 22 days after irradiation. Eggs laid by irradiated adults were collected daily, and the eggs were transferred with 90 µL of sterilized Milli-Q water to 96 well plates, whose each well was filled with 100 µL of 1.5% agar in the bottom. The hatchability of eggs was examined daily for approximately 3 weeks.

## RESULTS

### Effects of gamma radiation on the activity of adults

To evaluate the radiation tolerance of desiccation-sensitive tardigrade *G. myrops*, we exposed adult tardigrades to various doses of gamma radiation and examined the activity of the irradiated individuals at 24 hours or 48 hours after irradiation. For comparison, we also examined the radiotolerability of highly desiccation-tolerant species, *R. varieornatus* simultaneously in a similar manner. In both species, the proportion of active animals decreased as the irradiation dose was raised (Fig. 1). In *G. myrops*, the proportion of active animals at 48 hours after irradiation was 79 ± 10% at 2,000 Gy exposure and drastically dropped to 5.0 ± 8.5% at 3,000 Gy, ending up with no survival at 4,000 Gy. The median lethal dose (LD_50_) in 48 hours post-irradiation (LD_50/48h_) was calculated as 2,240 Gy (Table 1). In contrast, the proportion of active individuals in *R. varieornatus* were 81 ± 9.5% even at 5,000 Gy and decreased to 19 ± 8.5% at 7,000 Gy (Fig. 2). Its LD_50/48h_ was calculated as 5,970 Gy (Table 1). The LD_50/48h_ value of *G. myrops* was less than the half of that of *R. varieornatus*, indicating that desiccation-sensitive *G. myrops* is more sensitive to irradiation than highly desiccation-tolerant *R. varieornatus*. Similar trend was also observed at 24 hours post-irradiation (Fig. 1, Table 1).

**Figure 1.**
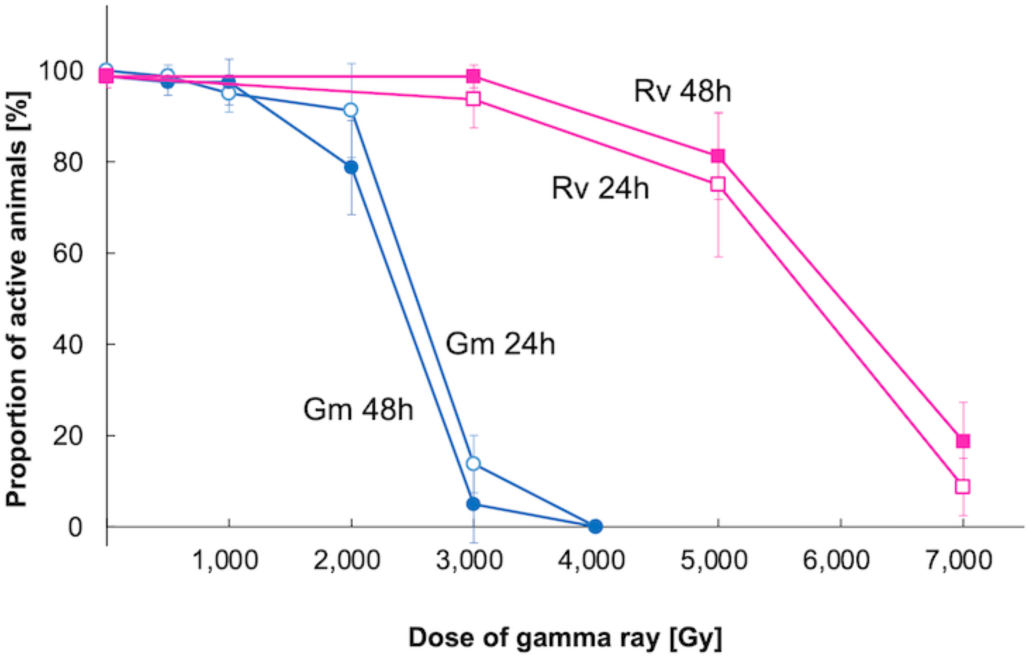
Effects of gamma irradiation on the activity of irradiated individuals in *Grevenius myrops* and *Ramazzottius varieornatus*. At 24 hours (open shapes) and 48 hours (closed shapes) after irradiation, the number of active animals were counted and the proportion was calculated. Squares and circles indicate the data for G*. myrops* (Gm) and *R. varieornatus* (Rv) respectively. Mean ± SD (*N* = 4; 19-20 tardigrades each).

**Figure 2.**
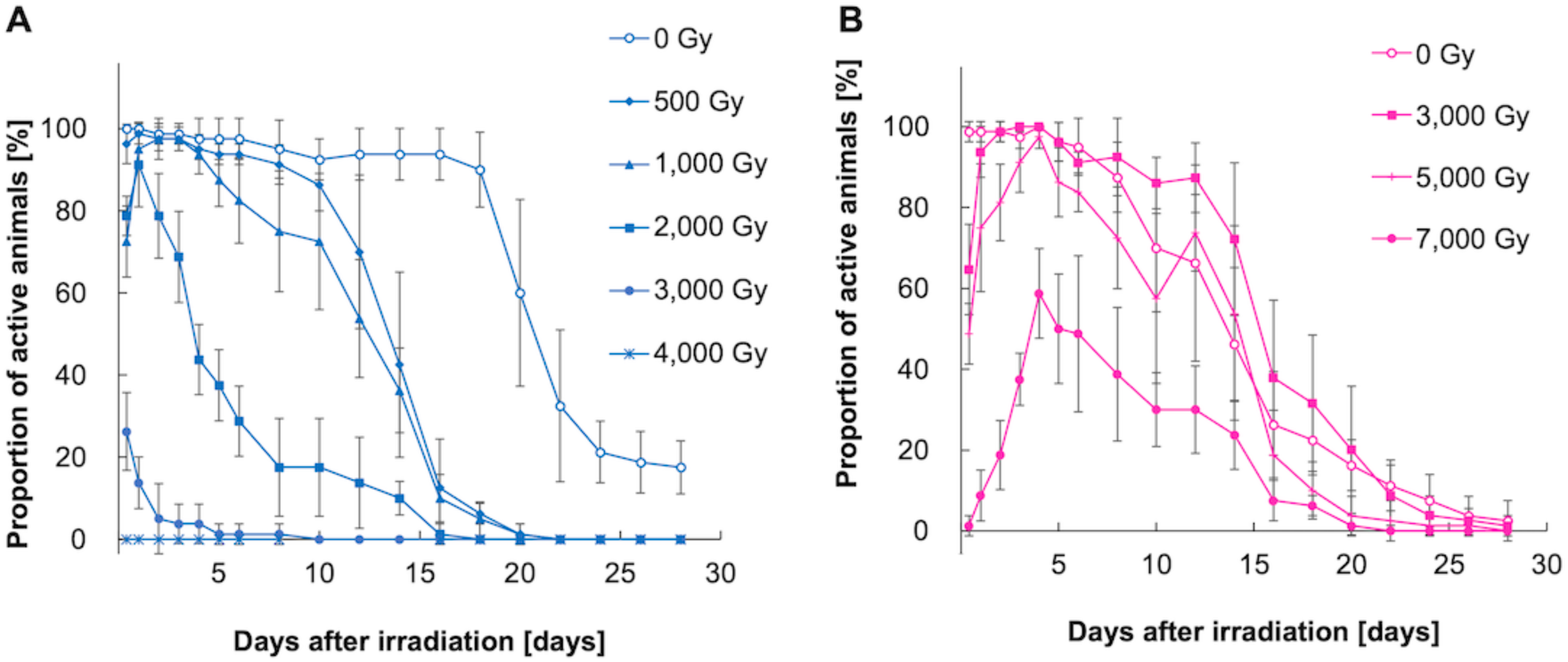
Temporal change of the proportion of active animals after gamma-irradiation. The activity of animals irradiated with various doses of gamma-ray were examined almost daily till 28 days after irradiation in *Grevenius myrops* (A) and *Ramazzottius varieornatus* (B). Mean ± SD (*N* = 4; 19-20 tardigrades each).

**Table 1.**
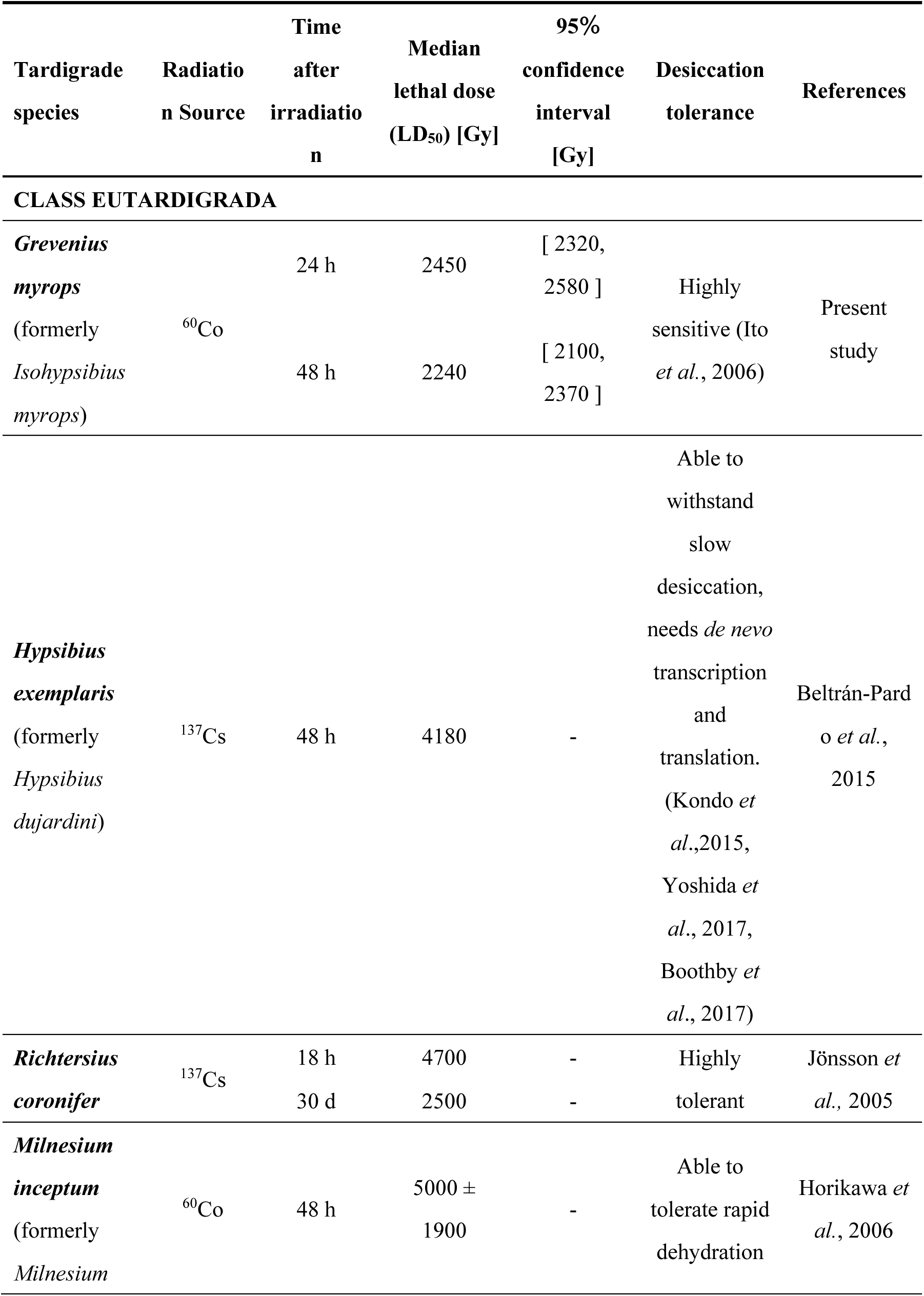
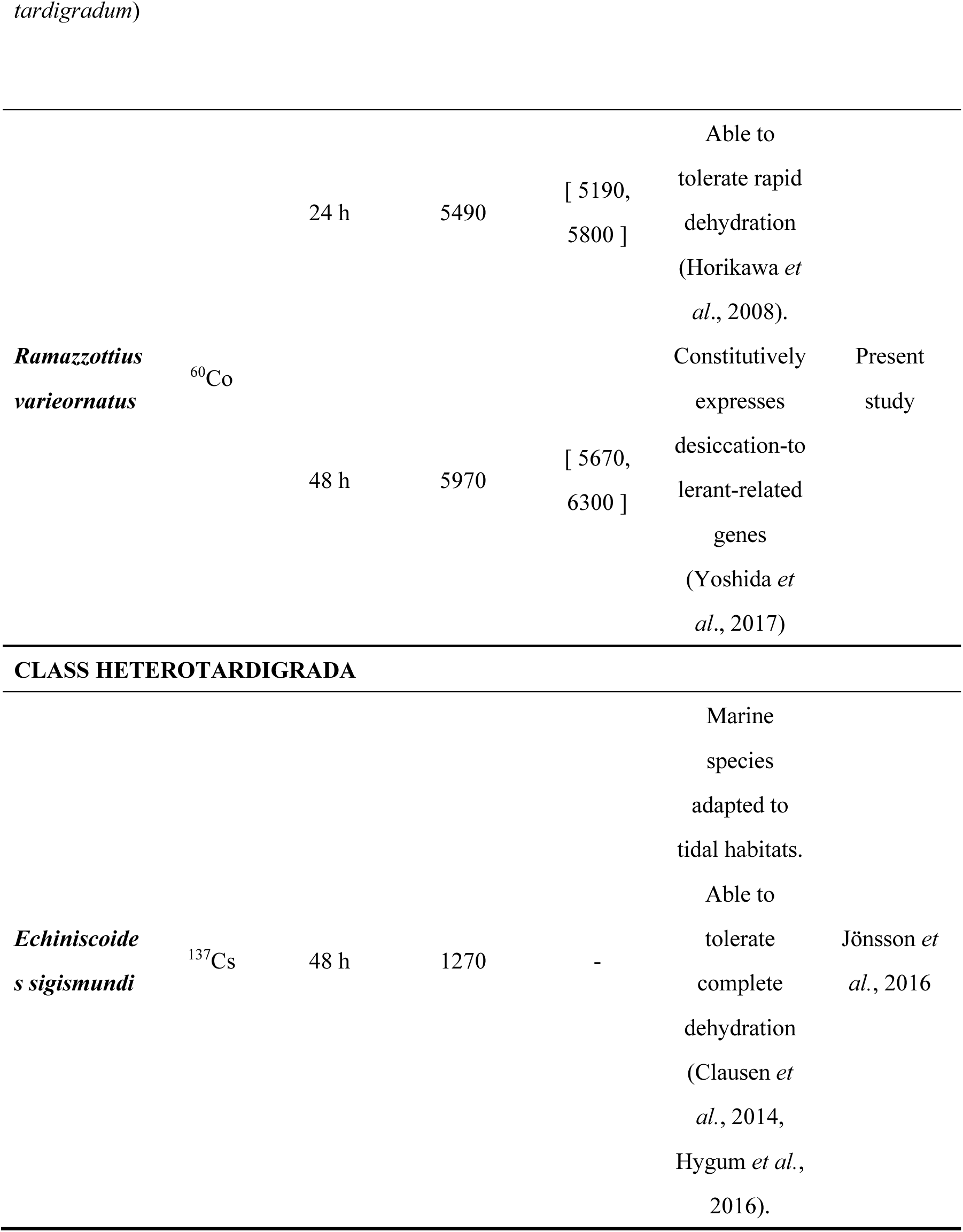
Tolerant ability of tardigrades against gamma irradiation in a hydrated state.

### Effects of gama radiation on lifespan

We next examined the effects of irradiation on the lifespan of adult tardigrades by continuous evaluation of the proportion of active individuals for 28 days almost daily. In *G. myrops*, shortly after irradiation (24 hours or 48 hours), a significant drop in the proportion of active individuals were observed only by irradiation of 3,000 Gy or higher doses, and with irradiation of 2,000 Gy or lower doses, the decreases were not so evident (Fig. 1). However, the continuous observations revealed that tardigrades irradiated with lower doses, e.g., even 500 Gy, exhibited significant reduction of the lifespan in *G. myrops* (Fig. 2A, Table 2). The median lifespan post-irradiation (i.e., the days after irradiation when 50% of animals ceased their activity) was approximately 21 days for 0 Gy (non-irradiated), whereas it was reduced to approximately 11–13 days by irradiation with 500 and 1,000 Gy. Irradiation with 2,000 Gy shortened the lifespan to approximately 5 days, which is almost 25% of the non-irradiated individuals. In contrast, the median lifespan of *R. varieornatus* was not significantly affected even by high dose (3,000–5,000 Gy) irradiation as the median lifespan was calculated as approximately 14 days at 0 Gy, and around 12–16 days at 3,000 or 5,000 Gy (Fig. 2B, Table 2).

**Table 2.**
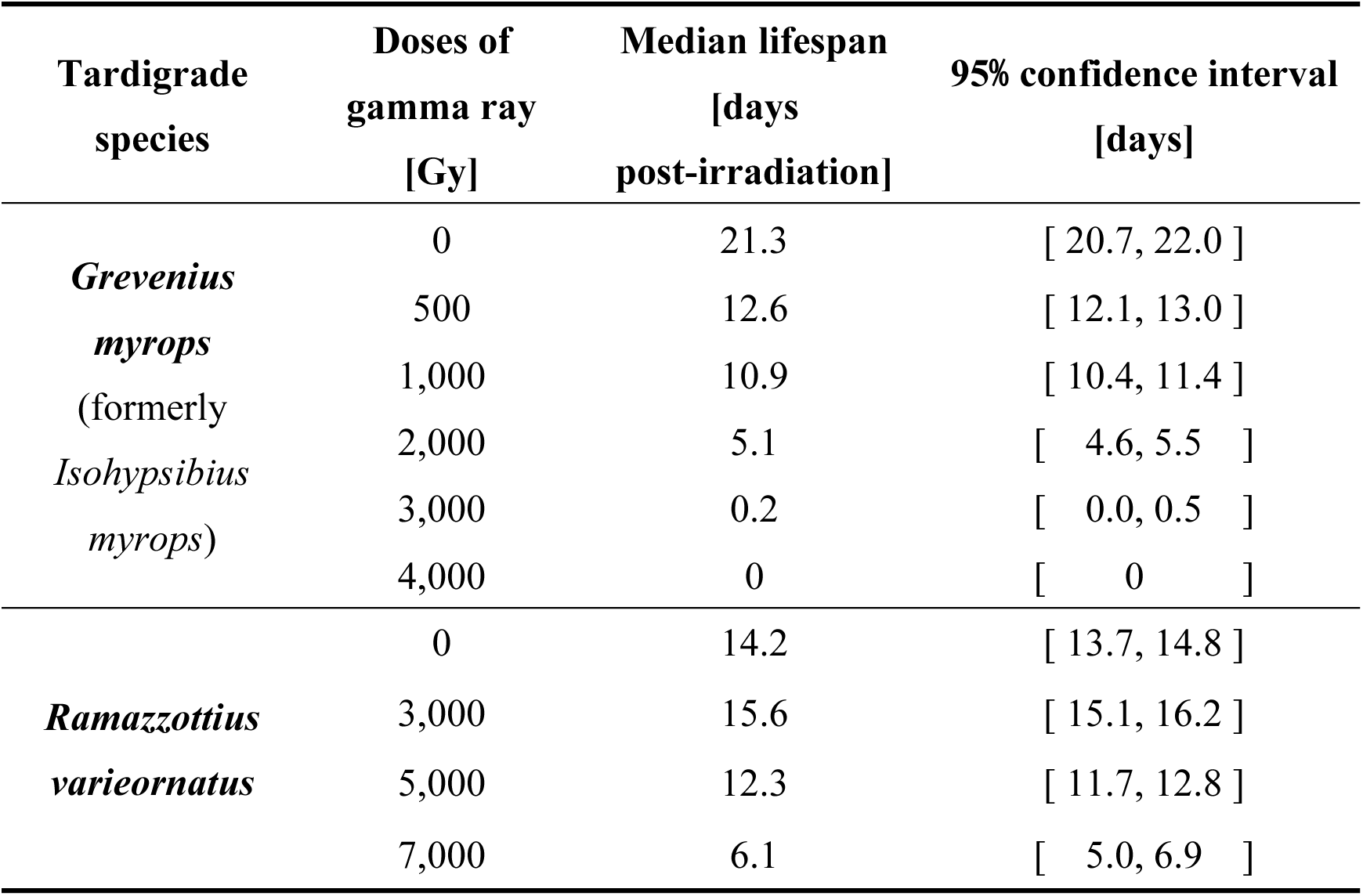
Effect of gamma radiation on the longevity of animals.

| **Tardigrade species** | **Doses of gamma ray [Gy]** | **Median lifespan [days post-irradiation]** | **95% confidence interval [days]** |
| --- | --- | --- | --- |
| ***Grevenius myrops***<br>(formerly *Isohypsibius myrops*) | 0 | 21.3 | [ 20.7, 22.0 ] |
|  | 500 | 12.6 | [ 12.1, 13.0 ] |
|  | 1,000 | 10.9 | [ 10.4, 11.4 ] |
|  | 2,000 | 5.1 | [ 4.6, 5.5 ] |
|  | 3,000 | 0.2 | [ 0.0, 0.5 ] |
|  | 4,000 | 0 | [ 0 ] |
| ***Ramazzottius varieornatus*** | 0 | 14.2 | [ 13.7, 14.8 ] |
|  | 3,000 | 15.6 | [ 15.1, 16.2 ] |
|  | 5,000 | 12.3 | [ 11.7, 12.8 ] |
|  | 7,000 | 6.1 | [ 5.0, 6.9 ] |

We also noticed a curious temporal change of the proportion of active animals after gamma irradiation. In *R. varieorantus* irradiated with 7,000 Gy, the proportion of active individuals temporarily dropped to 1.3 ± 2.5% at 10 hours, but then gradually increased to 59 ± 11% in 4 days after irradiation (Fig. 2B). Likewise, with 5,000 Gy irradiation, the proportion of the active individual was 49 ± 7.5% at 10 hours post-irradiation and then increased up to 98 ± 2.9% in 4 days post-irradiation. In *G. myrops*, individulas irradiated with 1,000 Gy or 2,000 Gy exhibited similar but slight increase in the proportion of active animals between 10 hours and 24 hours post-irradiation: i.e., from 73 ± 8.7% to 95 ± 4.1% with 1,000 Gy and from 79 ± 4.8% to 91 ± 10% with 2,000 Gy (Fig. 2A). However, these recoveries were not observed in *G. myrops* irradiated at 3,000 Gy. In *R. varieornatus*, most individuals became inactive with rolling up their bodies immediately after irradiation, but gradually began to move their legs coordinately, and in successful cases, they started to walk around on the agar just like the unirradiated individuals.

### Effects of gamma radiation on egg laying and hatchability

The previous observations suggested that the effect of gamma irradiation is more severe in germ line cells and early embryonic development (Jönsson *et al*., 2005; Horikawa *et al*., 2006; Beltrán-Pardo *et al*., 2015). Therefore, we further examined the effects of irradiation on egg laying and the hatchability of laid eggs. Both *G. myrops* and *R. varieornatus* propagate by parthenogenesis (Ito *et al*. 2016, Horikawa *et al*. 2008) and can reproduce offsprings without mating. Continuous observation for 22 days after irradiation revealed that the average number of eggs laid by each individual in *G. myrops* declined drastically even by low dose irradiation; i.e., from 45.3 ± 7.6 eggs at 0 Gy to 0.7 ± 0.3 eggs at 500 Gy and 0.3 ± 0.0 eggs at 1,000 Gy (Fig. 3A). Individuals irradiated with 2,000 Gy or higher doses did not lay any eggs. Some of the eggs produced by adults irradiated with 500 Gy exhibited deformed shapes such as inordinately small or ellipsoid shapes. In addition to these deformed shapes, with 1,000 Gy irradiation, we observed more frequently flat and wrinkle shaped eggs which seem to have just their eggshells. In *R. varieornatus*, the average number of laid eggs per individual was 3.9 ± 0.3 eggs at 0 Gy and decreased to 0.9 ± 0.3 eggs with 3,000 Gy (Fig. 3B). Individuals irradiated with 5,000 Gy or higher did not produce any eggs. Some of the eggs laid by adults irradiated with 3,000 Gy was much smaller than those of the non-irradiated group. In both species, all eggs laid by the irradiated adults failed to hatch (Fig. 3C, D).

**Figure 3.**
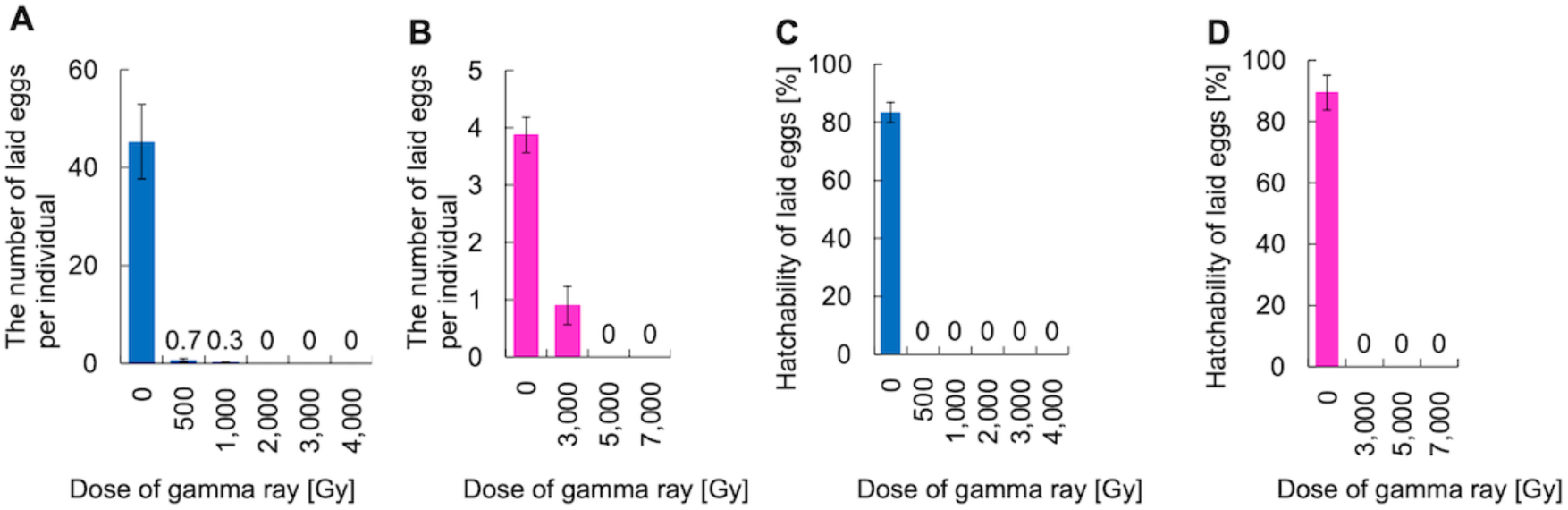
Effects of gamma radiation on oviposition in *Grevenius myrops* and *Ramazzottius varieornatus*. (A, B) The average number of laid eggs per individual in each group irradiated with various doses of gamma rays. Mean ± SD (*N* = 4; 19-20 tardigrades each). (C, D) The hatchability of eggs laid by adults irradiated with each dose of gamma-ray. (A, C) *Grevenius myrops*, (B, D) *Ramazzottius varieornatus*.

### Effects of gamma radiation on oviposition frequency

To reveal the temporal changes of oviposition by irradiated adults, the number of eggs laid by each group were counted for 23 days almost daily. Moving average with a window of 3 days was also plotted in order to visualize the long-term trends of oviposition frequency of each group (Fig. 4).

**Figure 4.**
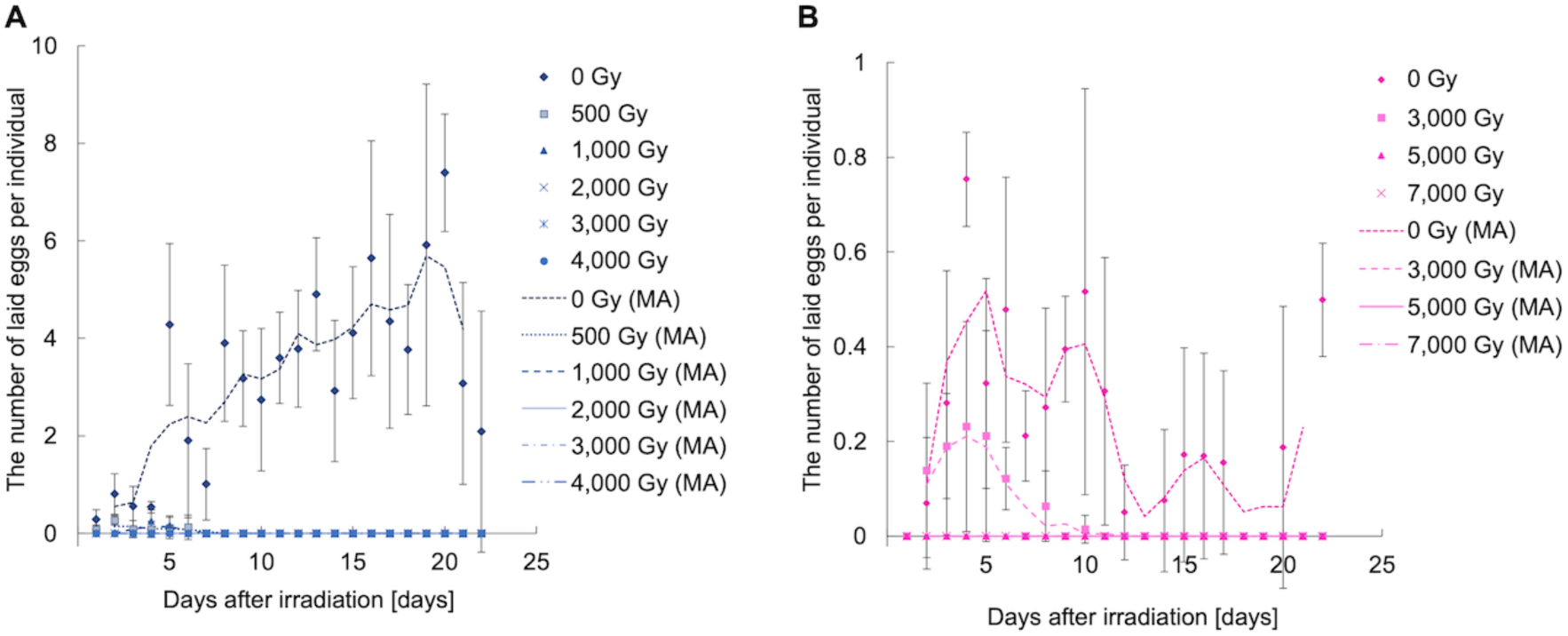
Temporal changes of the number of eggs laid by irradiated adults. The number of eggs laid by animals irradiated with various doses were examined almost daily. The moving average plots (labeled as “MA”) with a window of 3 days are also shown as lines in order to visualize the long-term trends of oviposition in each condition. (A) *Grevenius myrops,* (B) *Ramazzottius varieornatus*. Mean ± SD (N = 4; 19-20 tardigrades each).

In *G. myrops*, unirradiated adults laid eggs continuously and the number of laid eggs gradually increased as the adults grow (Fig. 4A). In contrast, irradiated individuals laid much fewer eggs only in a short period after irradiation; e.g., 6 days in 500 Gy groups and 5 days in 1,000 Gy groups. In individuals irradiated with 1,000 Gy or higher, we frequently observed the collapse and clouded structure of the ovary immediately after irradiation. In irradiated *R. varieornatus*, only one individual irradiated with 3,000 Gy did lay eggs till 10 days after irradiation (Fig 4B). All other irradiated individuals did not produce eggs.

## DISCUSSION

In this study, we provided the first data on the radiotolerability of a desiccation-sensitive tardigrade in details, including the effects on the activity and lifespan of the irradiated individuals and the effects on the oviposition and the hatchability. The median lethal dose (LD_50_) in 48 hours post-irradiation (LD_50/48h_) was calculated as approximately 2,240 Gy for desiccation-sensitive *G. myrops* and approximately 5,970 Gy for highly desiccation-tolerant *R. varieornatus* (Table 1, Fig. 1). In the literatures, two good desiccation-tolerant tardigrades species, *Richtersius coronifer* (Jönsson *et al*., 2005) and *Milnesium inceptum* (Horikawa *et al*., 2006) exhibited high LD_50_, around 4,700–5,000 Gy which are closer to that of *R. varieornatus* in this study (Table 1). Another tardigrade species, *Hypsibius exemplaris* (formerly *H. dujardini*; Gasiorek *et al*. 2018), which can tolerate only slow desiccation (Kondo *et al*., 2015, Yoshida *et al*., 2017) is reported to have a value of LD_50/48h_ of 4,180 Gy in a hydrated state (Beltran-Pardo *et al*., 2015), which corresponds around a medium value between those of *G. myrops* and *R. varieornatus* (Table 1). Apparently, there is the same trend in desiccation tolerance and radiotolerance among 5 tardigrade species which belong to class Eutardigrada and the data of both desiccation and radiation tolerance are available, *i.e.*, *G. myrops, R. varieornatus, R. coronifer, M. tardigradum* and *H. exemplaris*. These data aligned with the hypothesis that radiotolerance could be derived from desiccation tolerance and might share some mechanisms between two tolerances. On the other hand, in a marine tardigrade *Echiniscoides sigismundi* which is a good desiccation tolerant species belonging to another class Heterotardigrada, the value of LD_50/48h_ was reported as 1,270 Gy (Jönsson *et al*., 2016), much lower than that of *G. myrops.* Recent genome/transcriptome studies revealed that many desiccation tolerance genes were not shared between class Eutardigrada and class Heterotardigrada, suggesting that these two classes have adapted to terrestrial environments independently and have developed distinct molecular mechanisms coping with desiccation (Kamilari *et al*., 2019; Murai e*t al*., 2021; Yoshida and Tanaka 2022; Fleming *et al*., 2024). This could explain why *E. sigismundi* is an outlier in the trend found in 5 eutardigrade species.

Still, the value of LD_50_ in *G. myrops*, 2,240 Gy, can be regarded as relatively high. However, *C. elegans* is reported to maintain its survival for more than 5 days after exposure to ^137^Cs irradiation of 1,800 Gy (72-hour-young adult of the wild-type strain N2: Johnson and Hartman, 1988). In the case of freshwater monogonont rotifers, most of which are sensitive to desiccation, *Brachionus koreanus* is reported to have the LD_50/24h_ value of 2,900 Gy (Won *et al*., 2016). Taking these into consideration, the LD_50_ of *G. myrops* might not be classified in a high radiotolerance group at least among minute freshwater animals.

It should be noted that there is a caveat on the direct comparison across studies including ours, since there are many differences in the experimental conditions. First, in addition to the dose amount, the dose rates of irradiation varied among studies. Secondary, the source of radiation (*e.g.*, ^60^Co or ^137^Cs) might also affect the results. This was pointed out in *C. elegans* which were more sensitive to ^137^Cs than to ^60^Co (Hartman *et al*, 1996). Thirdly, specifically in the case of *E. sigismundi,* this species is a marine tardigrade and it was kept in seawater during exposure to radiation (Jönsson *et al*., 2016), whereas other eutardigrades were irradiated in freshwater. As the authors have already mentioned in their report of *E. sigismundi*, the difference of water might affect the results. Consequently, a concreate evidence will be obtained by a simultaneous comparison in the same experimental condition among various tardigrade species and other aquatic microorganisms in future.

In both tardigrade species, the egg laying was much more sensitively affected by radiation than the activity and the lifespan of adults (Fig. 3). Although some individuals laid a few eggs after relatively low dose irradiation, i.e., 500 Gy and 1000 Gy in *G. myrops* and 3,000 Gy in *R. varieornatus*, the ovipositions were limited to a short duration after irradiation and no adults were observed to lay eggs after 6–10 days post-irradiation (Fig. 4). These results implied that irradiated adults were able to lay eggs which were already produced before exposure to radiation, but failed to form new ones, therefore could not lay eggs from then on. In addition, all eggs laid by irradiated adults failed to hatch (Fig. 3C, D). These results are in a good agreement with the previous report that reproduction and/or germ cells are more sensitive to radiation than somatic cells (Jönsson *et al*., 2005; Horikawa *et al*., 2006; Beltrán-Pardo *et al*., 2015). We did not exclude a possibility that the gamma radiation impaired the health of the adults and repressed its oviposition or egg production.

In *R. varieornatus*, the average lifespan was virtually unaffected even after 5,000 Gy irradiation, which is a dose close to its LD_50/24h_, 5,490Gy (Table 2, Fig. 2B). In contrast, *G. myrops* exhibited a significantly shortened lifespan by radiation even with 500 Gy, which is much lower than its LD_50/24h_ (2,450 Gy) (Fig. 2A). This might indicate that low dose irradiation caused internal damage in a body of *G. myrops* which was invisible shortly after irradiation but shortened its lifespan. *G. myrops* may suffer more damage than *R. varieornatus* by low dose irradiation and/or have less ability to repair its internal damage. On the other hand, in *R. varieornatus*, especially clear with 5,000 Gy and 7,000 Gy irradiation, the proportion of active adults has temporarily dropped after irradiation but recovered during the following 4 days (Fig. 2B). This might suggest that *R. varieornatus* has an ability to repair its internal damage persistently at least for 4 days after irradiation, and this might be the reason why no significant decrease in the life span was observed in this species even after exposed to the dose near to its LD_50_. A similar phenomenon was reported in *R. coronifer* (Jönsson *et al*., 2005), which has relatively high desiccation tolerance and can withstand relatively fast dehydration. Nonetheless, in *M. tardigradum*, whose desiccation tolerance is as high as *R. varieornatus*, or in *H. exemplaris*, which can endure only slow dehydration, the recovery in the early phase was not observed and the proportion of active animal monotonically declined after irradiation as the time past (Horikawa *et al*., 2006, Beltran-Pardo *et al*., 2015). These results suggested that only limited desiccation-tolerant species have the ability to recover from the initial inactivation of irradiated individuals. Comparative study between *R. varieornatus* and *G. myrops* would shed light on the molecular mechanisms of these repair systems.

Currently, the mechanisms supporting high radiation tolerance of tardigrades are not fully understood, but several factors have been identified as involved in the radiotolerance of tardigrades. For example, Dsup (Damage suppressor) is a tardigrade-unique protein associated with DNA/chromatin and enhanced the tolerance against radiation, oxidative stress, and other genotoxic stresses, when introduced to human cultured cells, fly, plants, yeast and even *in vitro,* likely through protection of DNA from stressors (Hashimoto *et al*., 2016; Chavez *et al*., 2019; Kirke *et al*., 2020; Ricci *et al*., 2021; Zarubin *et al*., 2023; Aguilar *et al*., 2023; Ye *et al*., 2023; Casino *et al*., 2024). The protein sequences of Dsup are highly diverged among species, but based on the synteny in the genome, its orthologs have been identified in three eutardigrade species, *R. varieornatus*, *H*. *exemplaris* and *Paramacrobiotus metropolitanus* (Hashimoto *et al*., 2016; Hashimoto and Kunieda, 2017; Chavez *et al*., 2019; Hara *et al*., 2021; Sugiura *et al*., 2024). Another tardigrade-unique protein named TDR1 was recently identified through transcriptomic analyses in several tardigrade species and were shown to reduce DNA damage by a genotoxic reagent in human cultured cells (Anoud *et al*., 2024). Recent study indicated that the conserved DNA repair machinery is also important in the radiation tolerance of the tardigrade, *H. exemplaris* (Clark-Hachtel *et al*., 2024). Quite recently, the technologies for genome-editing and transient transgenesis became available in tardigrades (Kumagai *et al*., 2022; Kondo *et al*., 2024; Tanaka *et al*., 2023). To see what happens on the desiccation tolerance when manipulating the radiation tolerance genes, or vice versa, should be the important next steps to clarify the cross-tolerance between radiation tolerance and desiccation tolerance, and will shed lights on the evolutionary history of extraordinary radiotolerance in tardigrades.

## ACKNOWLEDGEMENTS

This work was supported by JSPS KAKENHI Grant Numbers JP16H02951, JP16H01632, JP18H04969, JP20H04332, JP20K20580, and JP21H05279.

## COMPETEING INTERESTS

The authors declared no competing interests.

## AUTHOR CONTRIBUTIONS

TS and TK conceived and designed the research. TS and KS performed experiments. TS analyzed the data. TS and TK wrote the draft. All authors reviewed and edited the final manuscript.

---

**Figure S1.**
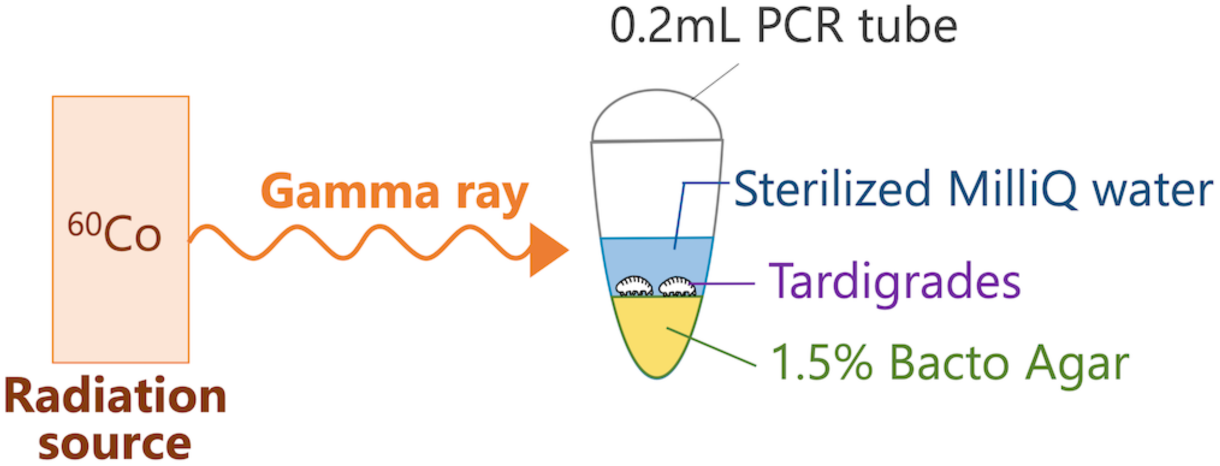
Schematic representation of the radiation apparatus.

## REFERENCES

1. Aguilar R, Khan L, Arslanovic N, Birmingham K, Kasliwal K, Posnikoff S, et al. (2023) Multivalent binding of the tardigrade Dsup protein to chromatin promotes yeast survival and longevity upon exposure to oxidative damage. Res Sq [Preprint] rs.3.rs-3182883

2. Anoud M, Delagoutte E, Helleu Q, Brion A, Duvernois-Berthet E, As M, et al. (2024) Comparative transcriptomics reveal a novel tardigrade specific DNA binding protein induced in response to ionizing radiation. eLife 13: RP92621

3. Beltrán-Pardo E, Jönsson KI, Harms-Ringdahl M, Haghdoost S, Wojcik A (2015) Tolerance to gamma radiation in the tardigrade Hypsibius dujardini from embryo to adult correlate inversely with cellular proliferation. PLoS ONE 10: e0133658

4. Billi D, Friedmann EI, Hofer KG, Caiola MG, Ocampo-Friedmann R (2000) Ionizing-radiation resistance in the desiccation-tolerant cyanobacterium Chroococcidiopsis. Appl Environ Microbiol 66(4): 1489–1492

5. Casino CD, Conti V, Licata S, Cai G, Cantore A, Ricci C, et al. (2024) Mitigation of UV-B radiation stress in tobacco pollen by expression of the tardigrade Damage suppressor protein (Dsup). Cells 13(10): 840

6. Chavez C, Cruz-Becerra G, Fei J, Kassavetis GA, Kadonaga JT (2019) The tardigrade damage suppressor protein binds to nucleosomes and protects DNA from hydroxyl radicals. eLife 8:e47682

7. Clark-Hachtel CM, Hibshman JD, De Buysscher T, Stair ER, Hicks LM, Goldstein B (2024) The tardigrade Hypsibius exemplaris dramatically upregulates DNA repair pathway genes in response to ionizing radiation. Curr Biol 34(9): 1819–1830.e6

8. Cox MM, Battista JR (2005) Deinococcus radiodurans – the consummate survivor. Nat Rev Microbiol 3(11): 882–892

9. Eutardigrada). Zootaxa 4415(1): 45–75

10. Fleming JF, Pisani D, Arakawa K (2024) The evolution of temperature and desiccation-related protein families in tardigrada reveals a complex acquisition of extremotolerance, Genome Biol Evol 16(1): evad217

11. Gąsiorek P, Stec D, Morek W, Michalczyk Ł (2018) An integrative redescription of Hypsibius dujardini (Doyère, 1840), the nominal taxon for Hypsibioidea (Tardigrada:

12. Gąsiorek P, Stec D, Morek W, Michalczyk Ł (2019) Deceptive conservatism of claws: distinct phyletic lineages concealed within Isohypsibioidea (Eutardigrada) revealed by molecular and morphological evidence. Contrib Zool 88: 78–132

13. Gladyshev EA, Meselson M, Arkhipova I (2008) Massive horizontal gene transfer in bdelloid rotifers. Science 320: 1210–1213

14. Hara Y, Shibahara R, Kondo K, Abe W, Kunieda T (2021) Parallel evolution of trehalose production machinery in anhydrobiotic animals via recurrent gene loss and horizontal transfer. Open Biol 11: 200413

15. Hartman P, Goldstein P, Algarra M, Hubbard D, Mabery J (1996) The nematode Caenorhabditis elegans is up to 39 times more sensitive to gamma radiation generated from 137Cs than from 60Co. Mutat Res 363(3):201–208

16. Hashimoto T, Horikawa D, Saito Y, Kuwahara H, Kozuka-Hata H, Shin-I T, et al. Extremotolerant tardigrade genome and improved radiotolerance of human cultured cells by tardigrade-unique protein. Nat Commun 7: 12808

17. Hashimoto T, Kuniedat T (2017) DNA protection protein, a novel mechanism of radiation tolerance: Lessons from tardigrades. Life (Basel) 7(2): 26

18. Horikawa DD, Kunieda T, Abe W, Watanabe M, Nakahara Y, Yukuhiro F, et al. (2008) Establishment of a rearing system of the extremotolerant tardigrade Ramazzottius varieornatus: A new model animal for astrobiology. Astrobiology 8: 549–556

19. Horikawa DD, Sakashita T, Katagiri C, Watanabe M, Kikawada T, Nakahara Y, et al. (2006) Radiation tolerance in the tardigrade Milnesium tardigradum. Int J Radiat Biol 82: 843–848

20. Ito M, Saigo T, Abe W, Kubo T, Kunieda T (2016) Establishment of an isogenic strain of the desiccation-sensitive tardigrade Isohypsibius myrops (Parachela, Eutardigrada) and its life history traits. Zool J Linn Soc 178(4): 863–870

21. Johnson TE, Hartman PS (1988) Radiation effects on life span in Caenorhabditis elegans. J Gerontol 43: B137–B141

22. Jönsson KI (2019) Radiation tolerance in tardigrades: Current knowledge and potential applications in medicine. Cancers 11(9): 1333

23. Jönsson KI, Harms-Ringdahl M, Torudd J (2005) Radiation tolerance in the eutardigrade Richtersius coronifer. Int J Radiat Biol 81: 649–656

24. Jönsson KI, Hygum TL, Andersen KN, Clausen LKB, Møbjerg N (2016) Tolerance to gamma radiation in the marine Heterotardigrade, Echiniscoides sigismundi. PLoS ONE 11: e0168884

25. Kamilari M, Jørgensen A, Schiøtt M, Møbjerg N (2019) Comparative transcriptomics suggest unique molecular adaptations within tardigrade lineages. BMC Genomics 20: 607

26. Kirke J, Jin X-L, Zhang X-H (2020) Expression of a tardigrade Dsup gene enhances genome protection in Plants. Mol Biotechnol 62(11-12): 563–571

27. Kondo K, Kubo T, Kunieda T (2015) Suggested involvement of PP1/PP2A activity and de novo gene expression in anhydrobiotic survival in a tardigrade, Hypsibius dujardini, by chemical genetic approach. PLoS ONE 10(12): e0144803

28. Kondo K, Tanaka A, Kunieda T (2024) Single-step generation of homozygous knockout/knock-in individuals in an extremototolerant parthenogenetic tardigrade using DIPA-CRISPR. PLOS Genet 20(6): e1011298

29. Kumagai H, Kondo K, Kunieda T (2022) Application of CRISPR/Cas9 system and the preferred no-indel end-joining repair in tardigrades. Biochem Biophys Res Commun 623: 196–201 Doi: 10.1016/j.bbrc.2022.07.060

30. Møbjerg N, Halberg KA, Jørgensen A, Persson D, Bjørn M, Ramløv H, et al. (2011) Survival in extreme environments – on the current knowledge of adaptations in tardigrades. Acta Physiol 202(3): 409–420

31. Murai Y, Yagi-Utsumi M, Fujiwara M, Tanaka S, Tomita M, Kato K, et al. (2021) Multiomics study of a heterotardigrade, Echinisicus testudo, suggests the possibility of convergent evolution of abundant heat-soluble proteins in Tardigrada. BMC Genomics 22: 813

32. Ricci C, Riolo G, Marzocchi C, Brunetti J, Pini A, Cantara S (2021) The tardigrade Damage suppressor protein modulates transcription factor and DNA repair genes in human cells treated with hydroxyl radicals and UV-C. Biology 10: 970

33. Ryabova A, Mukae K, Cherkasov A, Cornette R, Shagimardanova E, Sakashita T, et al. (2017) Genetic background of enhanced radioresistance in an anhydrobiotic insect: transcriptional response to ionizing radiations and desiccation. Extremophiles 21: 109–120

34. Sugiura K, Yoshida Y, Hayashi K, Arakawa K, Kunieda T, Matsumoto M (2024) Sexual dimorphism in the tardigrade Paramacrobiotus metropolitanus transcriptome. Zoological Lett 10: 11

35. Tanaka S, Aoki K, Arakawa K (2023) In vivo expression vector derived from anhydrobiotic tardigrade genome enables live imaging in Eutardigrada. Proc Natl Acad Sci USA 120(5): e2216739120

36. Won EJ, Han J, Hagiwara A, Oda S, Mitani H, Lee J-S (2016) Acute toxicity of gamma radiation to the monogonont rotifer Brachionus koreanus. Bull Environ Contam Toxicol 97: 387–391

37. Ye C, Guo J, Zhou X-Q, Chen D-G, Liu J, Peng X, et al. (2023) The Dsup coordinates grain development and abiotic stress in rice. Plant Physiol Biochem 205: 108184

38. Yoshida Y, Koutsovoulos G, Laetsch DR, Stevens L, Kumar S, Horikawa DD, et al. (2017) Comparative genomics of the tardigrades Hypsibius dujardini and Ramazzottius varieornatus. PLoS Biol 15(7): e2002266

39. Yoshida Y, Tanaka S (2022) Deciphering the biological enigma-genomic evolution underlying anhydrobiosis in the phylum Tardigrada and the chironomid Polypedilum vanderplanki. Insects 13(6):557

40. Zarubin M, Azorskaya T, Kuldoshina O, Alekseev S, Mitrofanov S, Kravchenko E (2023) The tardigrade Dsup protein enhances radioresistance in Drosophila melanogaster and acts as an unspecific repressor of transcription. iScience 26(7): 106998

